# Microbial associations with Microscopic Colitis

**DOI:** 10.1101/2021.12.10.472154

**Authors:** Shan Sun, Ivory C. Blakley, Anthony A. Fodor, Temitope O. Keku, John T. Woosley, Anne F. Peery, Robert S. Sandler

## Abstract

**BACKGROUND AND OBJECTIVE:** Microscopic colitis is a relatively common cause of chronic diarrhea and may be linked to luminal factors. Given the essential role of the microbiome in human gut health, analysis of microbiome changes associated with microscopic colitis could provide insights into the development of the disease.

**METHODS:** We enrolled patients who underwent colonoscopy for diarrhea. An experienced pathologist classified patients as having microscopic colitis (n=52) or controls (n=153). Research biopsies were taken from the ascending and descending colon, and the microbiome was characterized with Illumina sequencing. We analyzed the associations between microscopic colitis and microbiome with a series of increasingly complex models adjusted for a range of demographic and health factors.

**RESULTS:** We found that alpha-diversity was significantly lower in microscopic colitis cases compared to controls in the descending colon microbiome. In the descending colon, a series of models that adjusted for an increasing number of co-variates found taxa significantly associated with microscopic colitis, including Proteobacteria that was enriched in cases and *Collinsella* enriched in controls. While the alpha-diversity and taxa were not significantly associated with microscopic colitis in the ascending colon microbiome, the inference p-values based on ascending and descending microbiomes were highly correlated.

**CONCLUSION:** Our study demonstrates an altered microbiome in microscopic colitis cases compared to controls. Because both the cases and controls had diarrhea, we have identified candidate taxa that could be mechanistically responsible for the development of microscopic colitis independent of changes to the microbial community caused by diarrhea.

**Significance of this study:** *What is already known about this subject?:*

- Microscopic colitis is a common cause of chronic diarrhea. The exact etiology of microscopic colitis is unknown, but the gut microbiome is considered to play an important role.
- Prior studies have reported microbial changes associated with microscopic colitis, but the results are not consistent. Prior studies have generally involved small numbers of patients, fecal samples, and comparison to healthy controls.

*What are the new findings?:*

- The microbiome was altered in microscopic colitis patients.
- Microscopic colitis was associated with lower alpha-diversity, increase of inflammation related taxa and decrease of *Collinsella* in the descending colon microbiome independent of diarrhea status.
- The changes of microbial features were consistent between ascending and descending colon microbiomes, but with smaller changes in the ascending colon microbiome.

*How might it impact on clinical practice in the foreseeable future?:*

- The altered microbial communities in patients with microscopic colitis suggest that the patients might benefit from prevention or treatment with live biotherapeutic products.

## INTRODUCTION

Microscopic colitis is a common cause of chronic diarrhea ^1, 2^. The colon appears grossly normal during colonoscopy but there is a thickened collagen band or lymphocytic infiltration microscopically. Although microscopic colitis was previously thought to be rare, the incidence has increased in Europe and North America. ^3-9^ The incidence is comparable to Crohn’s disease and ulcerative colitis, conditions that have received much more research attention.

The exact etiology of microscopic colitis is unknown, but the gut microbiome is considered to play an important role. It is widely accepted that the condition represents an abnormal immune reaction to luminal antigens in the predisposed host.^10^ The hypothesis of the involvement of a luminal factor is supported by resolution of the disease with diversion of the fecal stream but recurrence when continuity is restored.^11, 12^ Fecal diversion also has a profound impact on the gut microbiome. ^13^ The human gut microbiome therefore likely plays important roles in the development of microscopic colitis. Investigation of the changes in microbiome composition associated with microscopic colitis could contribute to our understanding of the etiology of microscopic colitis and provide insights on treatment.

Prior studies^14-19^ on the microbiome associated with microscopic colitis have not always reported consistent results, and they have generally involved small numbers of patients, fecal samples, and comparison to healthy controls. Diarrhea in microscopic colitis patients can impact the microbial composition of their fecal samples ^20^, and this difference could contribute to the differences between the fecal samples of microscopic colitis patients and healthy controls. To learn more about the possible roles of bacteria in microscopic colitis, we conducted a prospective study at a single academic medical center. In this study, we recruited participants from patients who underwent colonoscopy for diarrhea. A research pathologist reviewed biopsies to determine whether the diarrhea was caused by microscopic colitis. We characterized the microbiomes of the colon biopsies with 16S rRNA sequencing and also systematically collected detailed demographic and exposure information from patient interviews.

## METHODS

### Study population and sample collection

We identified patients who were referred to the University of North Carolina Hospitals for diarrhea. Patients with known inflammatory bowel disease (IBD), *Clostridioides difficile* or infectious diarrhea based on chart review were excluded, as well as patients with gross evidence of IBD. Patients had to report loose stools as measured by the Bristol Stool Form Scale type 5-7. ^21^ Eligible patients signed informed consent, HIPPA and Storing Biological Specimens with Identifying Information form. During the colonoscopy, clinical biopsies were taken for standard pathologic review. Research biopsies were taken from the ascending colon and descending colon and immediately frozen in liquid nitrogen for later microbial analysis. Specimens were taken to the lab where they were stored at -80 °C. Patients were classified as microscopic colitis cases or controls by an experienced gastrointestinal pathologist (JTW). The microscopic colitis patients had increased numbers of intraepithelial or lamina propria lymphocytes or a thickened collagen band. The control group had neither increased lymphocytes nor thickened collage band. Patients with non-lymphocytic colitis were excluded. Patients were interviewed by phone about demographic factors, diet, medications, symptoms and autoimmune disease. Patients were asked if they had taken antibiotics in the three months prior to their colonoscopy. An identical interview was offered to some patients to be self-completed over the Internet.

### DNA extraction, PCR and Sequencing

Bacterial genomic DNA was extracted using previously described protocols. ^22, 23^ Normal colonic mucosal biopsies were placed in lysozyme containing buffer for 30 minutes followed by bead beating and DNA extraction with the Qiagen DNA Blood and Tissue Kit (Qiagen Cat# 69504). The purified DNA samples were stored in aliquots at −20□°C.

Illumina library preps were performed using previously published protocols. ^24, 25^ Briefly, PCR amplification was conducted in two separate reaction steps. The first PCR (PCR1) reaction contained the Phusion High-Fidelity Master Mix (Life Technologies, Carlsbad, CA) and primers targeting the V2 region of the 16S bacterial rRNA gene. The PCR1 product was diluted 20-fold and served as a template for the second PCR step (PCR2). PCR2 reaction mix contained primers with an Illumina index barcode sequence, Illumina adapter sequence, and a tag sequence. There were two sets of PCR2 primers. Each PCR2 reaction received one of each, resulting in a dual-indexed product. One reaction was performed for each sample using the Phusion High-Fidelity Master Mix. Using the E-Gel 96, we verified PCR amplification of samples. Those with positive amplification were normalized to 25 ng/µl (SequalPrep Normalization Kit; Life Technologies, Carlsbad, CA). An equal volume of each sample library was pooled followed by cleaning using AxyPrep Mag Beads. ^24^ The pool was stored at −20□°C, and later shipped to the University of Maryland Institute for Genome Sciences for Illumina MiSeq sequencing. ^24^ Positive and negative controls were included in all sample preparation steps. A pooled sample of known bacteria served as the positive control. The sequencing data analyzed in this study are available at NCBI as BioProject PRJNA768799.

### Sequence Processing and Statistical Analysis

The sequencing reads were analyzed with QIIME2 and DADA2 following the instructions. ^26, 27^ The forward reads were first truncated to 250 bp and denoised with DADA2 with the chimera removed using DADA2 ‘consensus’ method. The amplicon sequencing variants (ASVs) were then classified with the QIIME2 feature classifier classify-sklearn based on the SILVA database (release 132). ^28^ The taxonomic abundance tables were normalized as previously described to correct for the different sequencing depth across samples. ^29^ The statistical analyses and visualization were performed with R (version 4.0.5). The associations between the microbial community and case/control were analyzed with the PERMANOVA test using the R function ‘adonis’ in the package ‘vegan’. The principal coordinates analysis (PCoA) of the microbiomes were based on the Bray-Curtis dissimilarity at the genus level and visualized with functions in the same package. Shannon diversity was calculated with the function ‘diversity’ of the R package ‘vegan’ and used to characterize the alpha-diversity of the microbiome. We used multivariate adjusted linear regression models with the R function ‘lm’ to analyze the associations between case/control and Shannon diversity, PCoA coordinates and individual taxa. The covariates were selected based on their known associations with microscopic colitis or gut microbiome ^30-32^. We use four linear models: model 1 was adjusted for the covariates education, Proton Pump Inhibitors (PPIs) use and batch effects; model 2 included additional covariates sex and antibiotics use; model 3 was additionally adjusted for age; model 4 was additionally adjusted for body mass index (BMI). Rare taxa (prevalence <10% participants) were not included in order to avoid over adjustment for false discovery rate. P-values were adjusted for multiple hypotheses testing with the Benjamini-Hochberg method. Significant taxa were visualized on taxonomic trees using the function ‘tree_view’ in the R package ‘plotmicrobiome’ (https://github.com/ssun6/plotmicrobiome). Analysis scripts and metadata are publicly available at https://github.com/FodorLab/MicroscopicColitisMicrobiome.

### Patient and public involvement

The participants and public were not involved in the design or conduct of the study.

## RESULTS

The characteristics of study participants are shown in Table 1. The microscopic colitis cases were older than controls. Cases were more likely to be female and better educated. The BMI of cases was lower than controls. We characterized the ascending (ASC) and descending (DES) colon microbiomes by Illumina sequencing technology to determine whether the microbiomes were different between microscopic colitis cases and controls. We first analyzed the associations between case/control and the microbial community at the genus level with PCoA and univariate PERMANOVA tests. The case and control microbiomes were not separated at PCoA1 or PCoA2 for either ASC or DES microbiomes, but showed better separation at PCoA 3-6 (Fig. 1a-d and f-i). The PERMANOVA test indicated that the genus level composition was not significantly associated with case/control in the ASC microbiome (P=0.092), but was significant in the DES microbiome (R^2^=0.0087, P=0.043). We used four models that were adjusted for different variables to analyze the associations between Shannon diversity, PCoA 1-6 (Fig. 1e and j) and individual taxa (see methods). Model 1 was adjusted for education, Proton Pump Inhibitors (PPIs) use and batch effects; model 2 included additional covariates sex and antibiotics use; model 3 was additionally adjusted for age and model 4 was additionally adjusted for BMI. Shannon diversity of the ASC microbiomes was only significantly different between cases and controls for model 2 (Fig. 1d and e). However, in the DES microbiome, Shannon diversity was significantly lower in cases compared to controls in all four models (Fig. 1i and j). In the ASC microbiome, PCoA3 was significantly associated with microscopic colitis with all four models (Fig. 1e), and PCoA5 and PCoA6 were associated with microscopic colitis in some models (Fig. 1e). In the DES microbiome, PCoA5 was significantly associated with microscopic colitis in all four models (Fig. 1j).

**Table 1.**
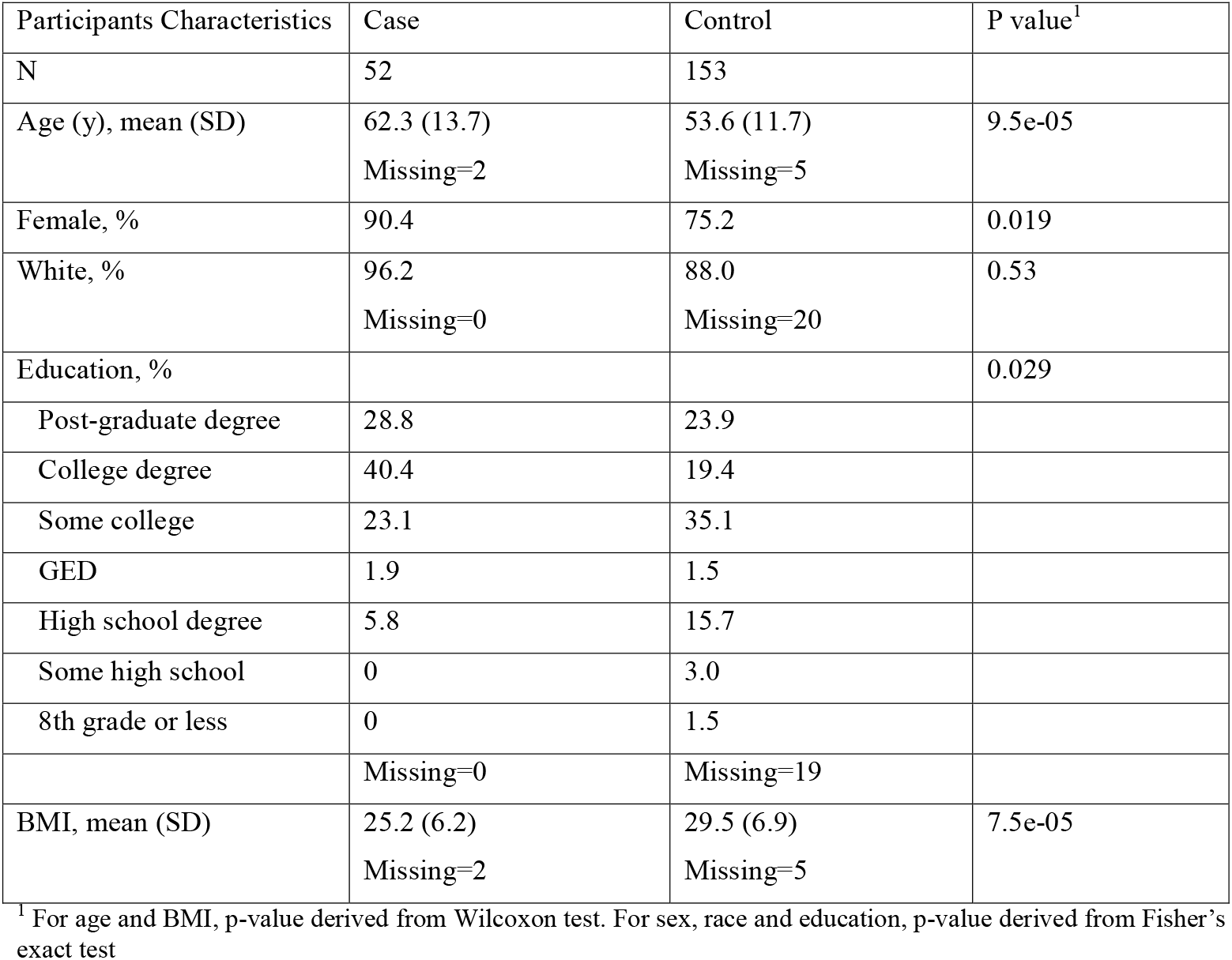
Characteristics of study participants.

**Fig. 1.**
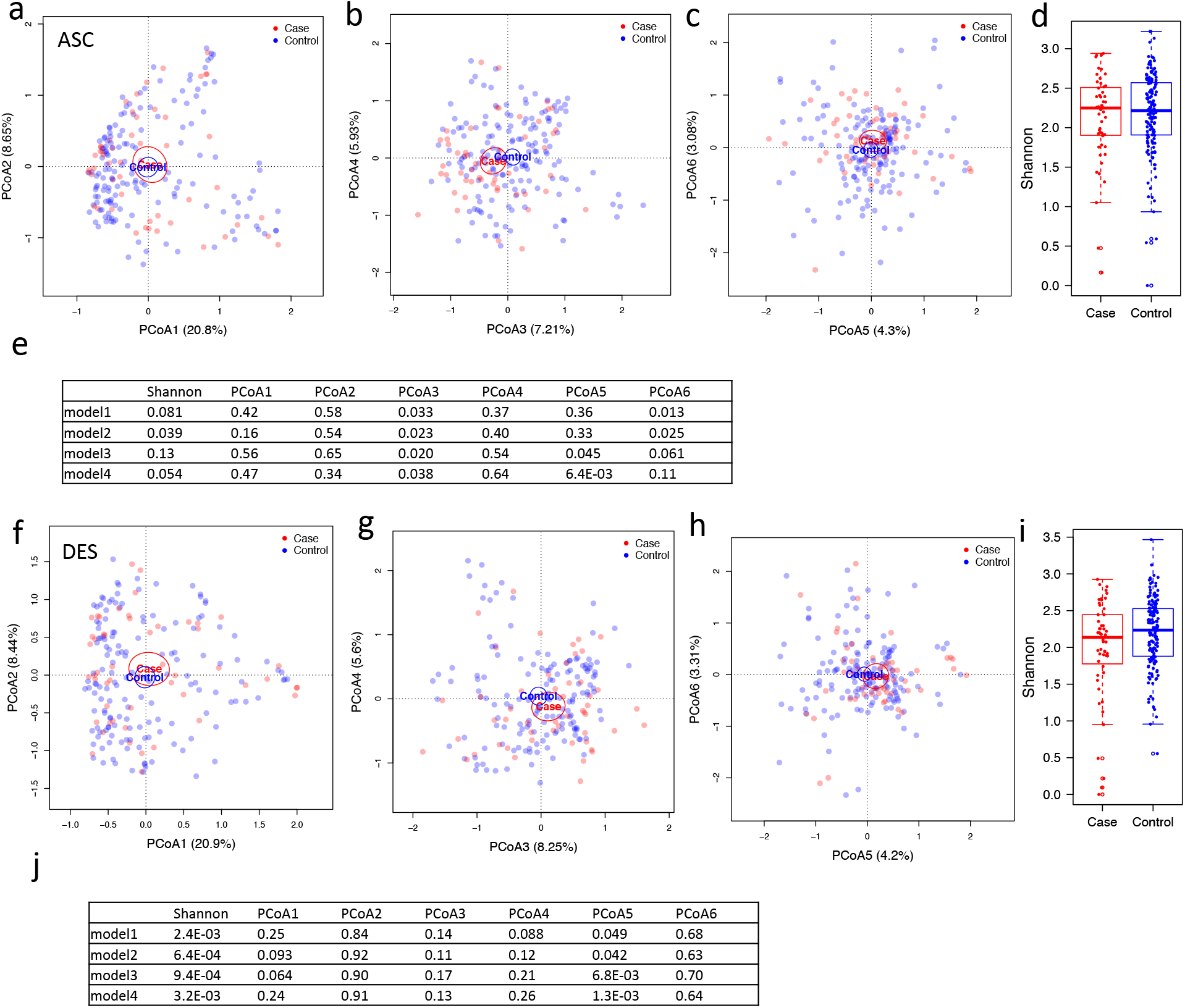
PCoA plots and alpha-diversity of the ascending (ASC) and descending (DES) colon microbiomes of study participants. (a-c) PCoA plots of the case and control microbiomes in the ASC microbiome showing PCoA 1-2 (a), 3-4 (b) and 5-6 (c). (d) Boxplots of the Shannon diversity of cases and controls in the ASC microbiome. (e) P-values of linear regression models analyzing the associations between case/control and Shannon diversity, PCoA1-6 adjusted for covariates (see methods) in the ASC microbiome. (f-h) PCoA plots of case and control microbiomes in the DES microbiome. (i) Boxplots of the Shannon diversity of cases and controls for the DES microbiome. (j) The P-values of linear regression models analyzing the associations between case/control and Shannon index, PCoA1-6 adjusted for covariates (see methods) in the DES microbiome. The ellipses in PCoA plots indicate 95% confidence limits. The boxplots showed the median, 25th and 75th percentile.

We also used the four models to analyze the associations between individual taxa and case/control to identify the differential taxa associated with microscopic colitis. The taxa of the ASC colon microbiome were not significantly associated with case/control in any of the models after adjusting for multiple testing, while there were significantly associated taxa in the DES microbiome (Table 2, FDR < 0.1). Model 1 (adjusted for batch, education and PPI) and model 2 (additionally adjusted for sex and antibiotics use) revealed similar differential taxa (21 in common, 3 only in model 1 and 6 only in model 2) (Fig. 2). However, model 3 revealed only 4 taxa after being additionally adjusted for age, model 4 revealed only 2 taxa after being additionally adjusted for BMI (Fig. 2). Because age and BMI were significantly different between case and control participants, it is possible that age and BMI confounded the microbiome associations with case/control.

**Table 2.** The P-values of linear regression models analyzing the associations between individual taxa and case/control adjusted for covariates (see methods) in the ascending (ASC) and descending (DES) colon microbiome

|  | ASCENDING COLON |  |  |  |  |  |  |  | DESCENDING COLON |  |  |  |  |  |  |  |
| --- | --- | --- | --- | --- | --- | --- | --- | --- | --- | --- | --- | --- | --- | --- | --- | --- |
|  | model1 |  | model2 |  | model3 |  | model4 |  | model1 |  | model2 |  | model3 |  | model4 |  |
|  | t-value | FDR | t-value | FDR | t-value | FDR | t-value | FDR | t-value | FDR | t-value | FDR | t-value | FDR | t-value | FDR |
| p__Proteobacteria--c__Gammaproteobacteria--o__Betaproteobacteriales | -3.22 | 0.26 | -3.11 | 0.18 | -3.35 | 0.26 | -3.36 | 0.23 | -4.31 | **0.009** | -4.14 | **0.017** | -4.18 | **0.015** | -3.36 | **0.011** |
| p__Proteobacteria--c__Gammaproteobacteria--o__Betaproteobacteriales--f__Burkholderiaceae | -2.96 | 0.26 | -2.89 | 0.2 | -3.2 | 0.26 | -3.24 | 0.23 | -3.87 | **0.024** | -3.76 | **0.037** | -3.77 | **0.036** | -3.24 | **0.027** |
| p__Proteobacteria--c__Gammaproteobacteria--o__Betaproteobacteriales--f__Burkholderiaceae--g__Sutterella--s__unclassified | -2.55 | 0.28 | -2.53 | 0.24 | -3 | 0.33 | -2.78 | 0.28 | -3.42 | **0.058** | -3.33 | **0.045** | -3.49 | **0.066** | -2.78 | 0.15 |
| p__Proteobacteria--c__Gammaproteobacteria--o__Pasteurellales--f__Pasteurellaceae--g__Haemophilus--s__unclassified | -1.04 | 0.79 | -1.08 | 0.69 | -1.01 | 0.93 | -0.62 | 0.94 | -3.38 | **0.058** | -3.18 | **0.045** | -2.99 | 0.11 | -0.62 | 0.23 |
| p__Proteobacteria--c__Gammaproteobacteria--o__Pasteurellales--f__Pasteurellaceae--g__Haemophilus | -1.04 | 0.79 | -1.08 | 0.69 | -1.01 | 0.93 | -0.62 | 0.94 | -3.38 | **0.058** | -3.18 | **0.045** | -2.98 | 0.11 | -0.62 | 0.23 |
| p__Proteobacteria--c__Gammaproteobacteria--o__Betaproteobacteriales--f__Burkholderiaceae--g__Sutterella | -2.31 | 0.35 | -2.32 | 0.31 | -2.75 | 0.37 | -2.54 | 0.38 | -3.2 | **0.063** | -3.16 | **0.045** | -3.35 | **0.079** | -2.54 | 0.17 |
| p__Proteobacteria--c__Gammaproteobacteria--o__Pasteurellales | -1.14 | 0.71 | -1.17 | 0.64 | -1.02 | 0.93 | -0.67 | 0.94 | -3.04 | **0.063** | -2.83 | **0.081** | -2.48 | 0.2 | -0.67 | 0.33 |
| p__Proteobacteria--c__Gammaproteobacteria--o__Pasteurellales--f__Pasteurellaceae | -1.14 | 0.71 | -1.17 | 0.64 | -1.02 | 0.93 | -0.67 | 0.94 | -3.04 | **0.063** | -2.83 | **0.081** | -2.48 | 0.2 | -0.67 | 0.33 |
| p__Proteobacteria--c__Gammaproteobacteria | -2.95 | 0.26 | -3.31 | 0.18 | -2.75 | 0.37 | -2.9 | 0.28 | -3.03 | **0.063** | -3.25 | **0.045** | -2.47 | 0.2 | -2.9 | 0.21 |
| p__Proteobacteria | -2.91 | 0.26 | -3.28 | 0.18 | -2.73 | 0.37 | -2.88 | 0.28 | -2.96 | **0.063** | -3.17 | **0.045** | -2.38 | 0.2 | -2.88 | 0.21 |
| p__Firmicutes--c__Bacilli--o__Lactobacillales--f__Streptococcaceae--g__Streptococcus--s__unclassified | -1.47 | 0.54 | -1.22 | 0.62 | -1.61 | 0.65 | -1.58 | 0.69 | -2.95 | **0.063** | -2.83 | **0.081** | -2.93 | 0.11 | -1.58 | 0.21 |
| p__Firmicutes--c__Bacilli--o__Lactobacillales | -1.62 | 0.52 | -1.39 | 0.56 | -1.7 | 0.63 | -1.72 | 0.65 | -2.94 | **0.063** | -2.83 | **0.081** | -2.87 | 0.11 | -1.72 | 0.21 |
| p__Firmicutes--c__Bacilli--o__Lactobacillales--f__Streptococcaceae | -1.46 | 0.54 | -1.22 | 0.62 | -1.62 | 0.65 | -1.59 | 0.69 | -2.93 | **0.063** | -2.82 | **0.081** | -2.91 | 0.11 | -1.59 | 0.21 |
| p__Firmicutes--c__Bacilli--o__Lactobacillales--f__Streptococcaceae--g__Streptococcus | -1.47 | 0.54 | -1.22 | 0.62 | -1.62 | 0.65 | -1.59 | 0.69 | -2.93 | **0.063** | -2.82 | **0.081** | -2.92 | 0.11 | -1.59 | 0.21 |
| p__Bacteroidetes--c__Bacteroidia--o__Bacteroidales--f__Marinifilaceae | -2.82 | 0.26 | -2.79 | 0.21 | -2.37 | 0.41 | -2.11 | 0.48 | -2.88 | **0.068** | -2.77 | **0.083** | -2.2 | 0.24 | -2.11 | 0.33 |
| p__Proteobacteria--c__Gammaproteobacteria--o__Betaproteobacteriales--f__Neisseriaceae--g__Neisseria | -1.96 | 0.4 | -1.68 | 0.51 | -1.23 | 0.86 | -1 | 0.9 | -2.74 | **0.094** | -2.42 | 0.16 | -2.49 | 0.2 | -1 | 0.21 |
| p__Proteobacteria--c__Gammaproteobacteria--o__Betaproteobacteriales--f__Neisseriaceae--g__Neisseria--s__unclassified | -1.96 | 0.4 | -1.68 | 0.51 | -1.23 | 0.86 | -1 | 0.9 | -2.74 | **0.094** | -2.42 | 0.16 | -2.49 | 0.2 | -1 | 0.21 |
| p__Proteobacteria--c__Gammaproteobacteria--o__Betaproteobacteriales--f__Neisseriaceae | -2.05 | 0.37 | -1.78 | 0.51 | -1.32 | 0.8 | -1.13 | 0.84 | -2.73 | **0.094** | -2.41 | 0.16 | -2.48 | 0.2 | -1.13 | 0.21 |
| p__Bacteroidetes--c__Bacteroidia--o__Bacteroidales--f__Marinifilaceae--g__Odoribacter--s__unclassified | -2.08 | 0.37 | -2.34 | 0.31 | -2.2 | 0.41 | -2.29 | 0.4 | -2.52 | 0.15 | -2.71 | **0.09** | -2.38 | 0.2 | -2.29 | 0.21 |
| p__Proteobacteria--c__Gammaproteobacteria--o__Enterobacteriales | -2.64 | 0.28 | -3.03 | 0.18 | -2.47 | 0.41 | -2.67 | 0.3 | -2.47 | 0.16 | -2.79 | **0.08** | -2.03 | 0.3 | -2.67 | 0.32 |
| p__Proteobacteria--c__Gammaproteobacteria--o__Enterobacteriales--f__Enterobacteriaceae | -2.64 | 0.28 | -3.03 | 0.18 | -2.47 | 0.41 | -2.67 | 0.3 | -2.47 | 0.16 | -2.79 | **0.08** | -2.03 | 0.3 | -2.67 | 0.32 |
| p__Firmicutes--c__Clostridia--o__Clostridiales--f__Lachnospiraceae--g__Blautia | 1.92 | 0.4 | 2.14 | 0.34 | 1.14 | 0.9 | 0.99 | 0.9 | 2.36 | 0.19 | 2.67 | **0.1** | 1.58 | 0.48 | 0.99 | 0.56 |
| p__Firmicutes--c__Clostridia--o__Clostridiales--f__Lachnospiraceae--g__Blautia--s__unclassified | 1.92 | 0.4 | 2.14 | 0.34 | 1.14 | 0.9 | 0.99 | 0.9 | 2.36 | 0.19 | 2.67 | **0.1** | 1.58 | 0.48 | 0.99 | 0.56 |
| p__Firmicutes--c__Negativicutes--o__Selenomonadales--f__Acidaminococcaceae--g__Phascolarctobacterium--s__unclassified | 2.09 | 0.37 | 2.47 | 0.26 | 2.58 | 0.41 | 2.37 | 0.4 | 2.62 | 0.12 | 2.86 | **0.08** | 2.88 | 0.11 | 2.37 | 0.21 |
| p__Actinobacteria | 2.19 | 0.36 | 2.32 | 0.31 | 1.95 | 0.46 | 1.68 | 0.66 | 2.88 | **0.07** | 3.05 | **0.06** | 2.38 | 0.2 | 1.68 | 0.3 |
| p__Actinobacteria--c__Coriobacteriia--o__Coriobacteriales--f__Coriobacteriaceae--g__Collinsella | 2.5 | 0.28 | 2.56 | 0.24 | 2.14 | 0.41 | 1.93 | 0.54 | 3.09 | **0.06** | 3.28 | **0.05** | 2.67 | 0.16 | 1.93 | 0.21 |
| p__Actinobacteria--c__Coriobacteriia--o__Coriobacteriales--f__Coriobacteriaceae--g__Collinsella--s__unclassified | 2.5 | 0.28 | 2.56 | 0.24 | 2.14 | 0.41 | 1.93 | 0.54 | 3.09 | **0.06** | 3.28 | **0.05** | 2.67 | 0.16 | 1.93 | 0.21 |
| p__Actinobacteria--c__Coriobacteriia--o__Coriobacteriales--f__Coriobacteriaceae | 2.53 | 0.28 | 2.58 | 0.24 | 2.16 | 0.41 | 1.95 | 0.54 | 3.1 | **0.06** | 3.3 | **0.05** | 2.69 | 0.16 | 1.95 | 0.21 |
| p__Actinobacteria--c__Coriobacteriia | 2.8 | 0.26 | 2.85 | 0.2 | 2.43 | 0.41 | 2.19 | 0.44 | 3.27 | **0.06** | 3.45 | **0.05** | 2.92 | 0.11 | 2.19 | 0.21 |
| p__Actinobacteria--c__Coriobacteriia--o__Coriobacteriales | 2.8 | 0.26 | 2.85 | 0.2 | 2.43 | 0.41 | 2.19 | 0.44 | 3.27 | **0.06** | 3.45 | **0.05** | 2.92 | 0.11 | 2.19 | 0.21 |

**Fig. 2.**
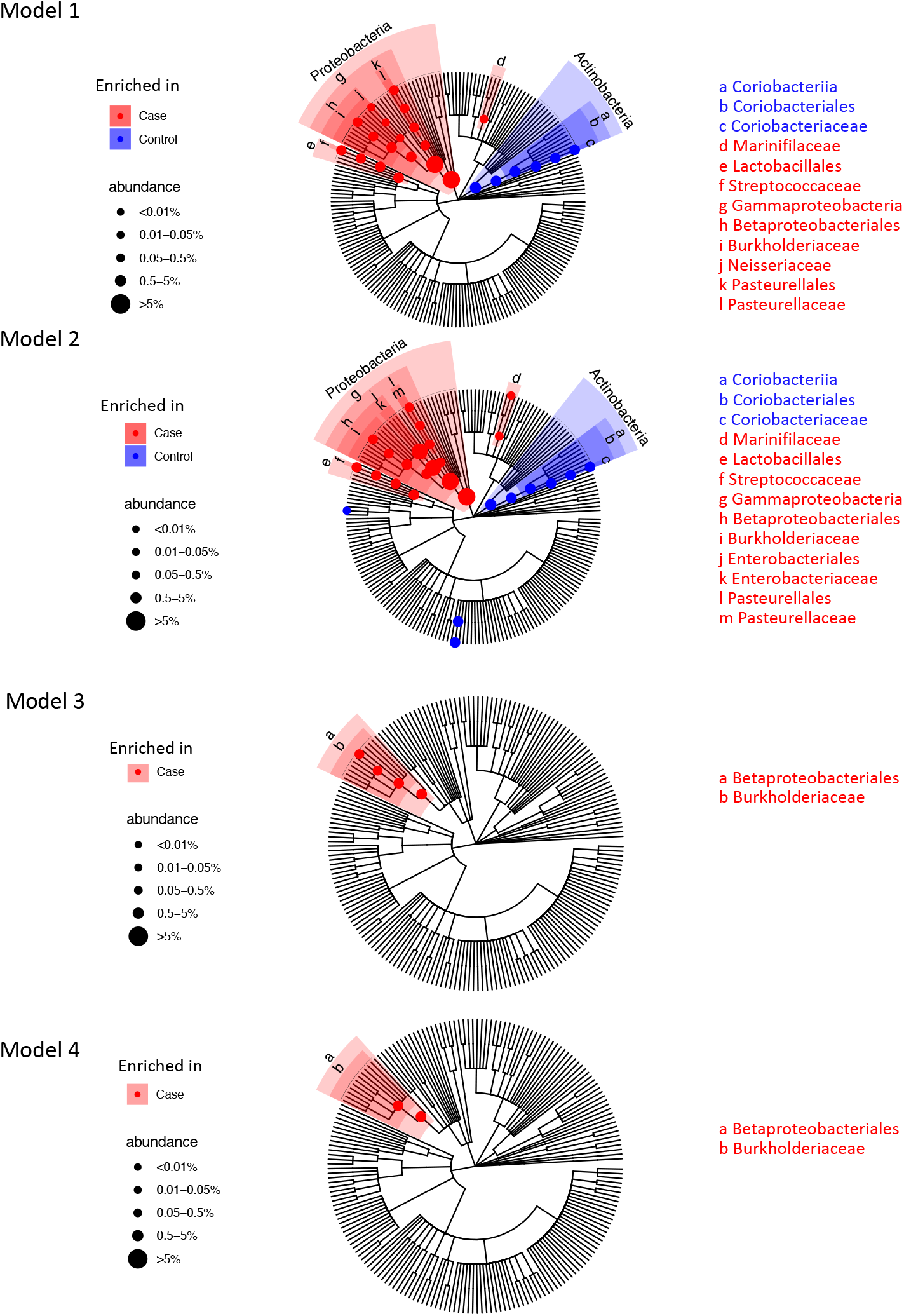
Significant differential taxa between microscopic colitis cases and controls highlighted in the taxonomic tree of the descending (DES) colon microbiome for the four linear regression models. Model 1 is adjusted for batch, education and PPI use. Model 2 is additionally adjusted for sex and antibiotics. Model 3 is adjusted for age along with the covariates in model 2. Model 4 is adjusted for BMI along with the covariates in model 3. The branches of significant taxa from phylum to family level were highlighted and labeled. The node sizes are proportional to the overall relative abundance of the taxa.

Among the 24 differential taxa revealed with model 1, 18 taxa were more abundant in cases and 6 were more abundant in controls, while among the 27 differential taxa revealed with model 2, 18 enriched in cases and 9 enriched in controls. The taxa enriched in the controls mostly belonged to Actinobacteria, mainly driven by genus *Collinsella* and its higher taxonomic levels (Fig 2 and Table 2). However, these associations were not significant in models that included age (model 3). The taxa enriched in cases were mostly Gammaproteobacteria, including *Sutterella, Haemophilus, Neisseria, Enterobacteriaceae* and *Pasteurellaceae*. Genus *Streptococcus* in phylum Firmicutes and family *Marinifilaceae* in phylum Bacteroidetes were also enriched in microscopic colitis cases (Fig 2 and Table 2). While the number of taxa associated with case decreased with increasingly model complexity, there were taxa associated with case in all four models suggesting that confounding with age and BMI cannot explain all of the associations we saw with microscopic colitis. The taxa revealed by model 3 were enriched in cases, including genus *Sutterella*, unclassified species in *Sutterella, Burkholderiaceae* and Betaproteobacteriales, while the taxa revealed by model 4 were *Burkholderiaceae* and Betaproteobacteriales that were enriched in cases. While we found no significant taxa associated with case/control in the ASC microbiome, the FDR corrected p-values estimated from the ASC and DES microbiomes were highly correlated (Fig. 3), indicating that while the microbial features associated with microscopic colitis were stronger in DES than ASC, a similar pattern of difference was seen at both sampling sites.

**Fig. 3.**
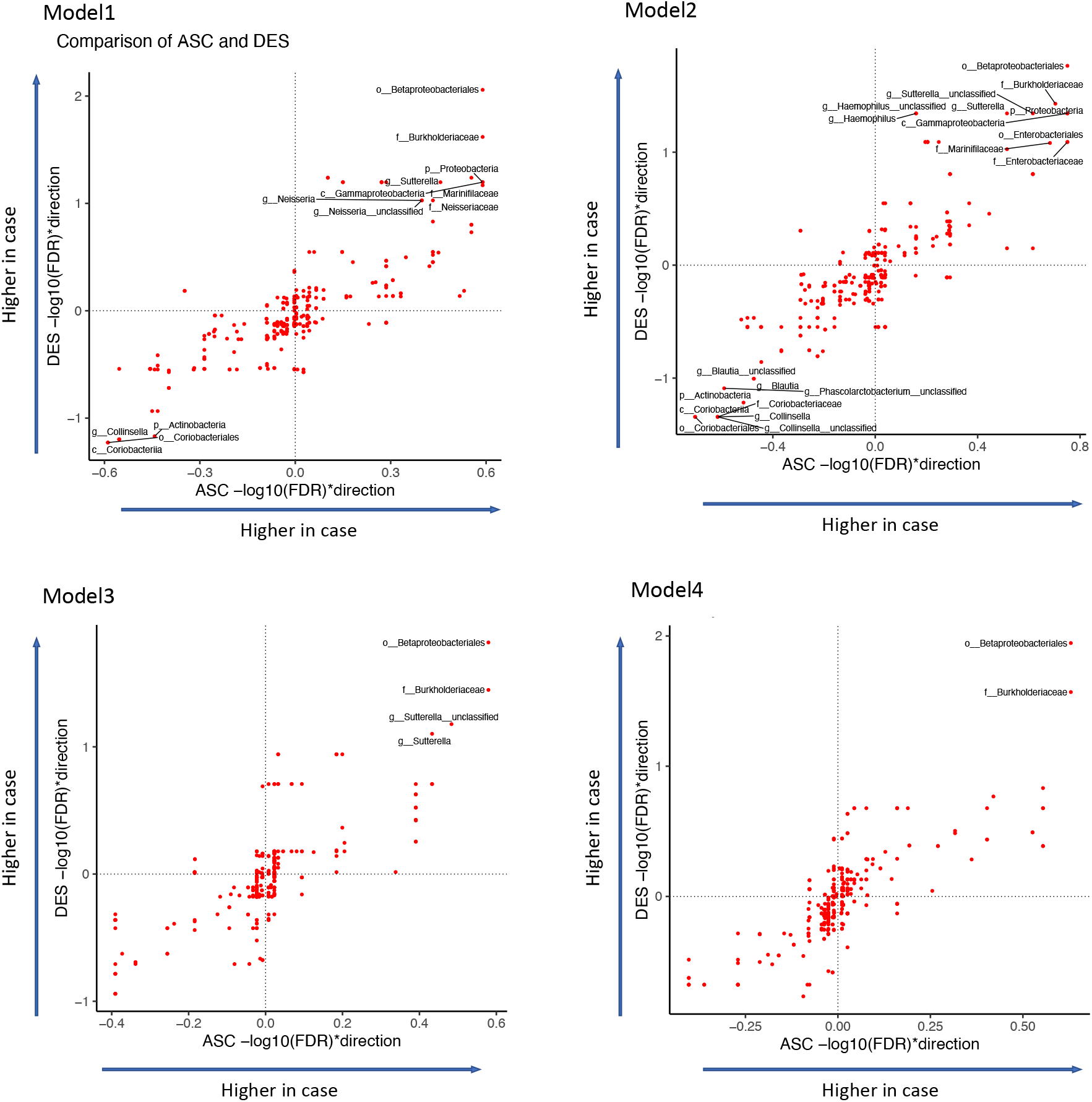
The comparison of the associations of individual taxa and case/control in ASC and DES microbiome with the four models. The x and y axes show the –log10 (FDR) multiplied by +1/-1 to indicate the direction of changes. Significant taxa (FDR <0.1) were labeled.

## DISCUSSION

In this study of the microbiome of microscopic colitis cases and diarrhea controls, we found that alpha diversity was significantly lower in cases than controls in the descending colon microbiome. We also found microorganisms that are associated with microscopic colitis, including taxa in phylum Proteobacteria that are potentially inflammation related. These differential taxa remained significant after adjusting the models for demographic factors and medicines, including sex, education, PPI and antibiotic use. Patient age and BMI were significantly associated with case/control status. Some taxa were not significant after adjusting for age and BMI. While we did not find significant differences in the ascending colon microbiome, the inference p-values were highly similar between the ascending and descending microbiome, indicating consistent but reduced microbial changes in the ascending microbiome compared to the descending microbiome.

While the etiology of microscopic colitis remains unknown, there are some clues. In 1995, Järnerot et al described the experience of 20 patients with microscopic colitis with severe diarrhea and increased thickness of the subepithelial collagen layer.^11^ Following fecal diversion, the diarrhea stopped in all patients and the collagen layer thinned. Symptoms recurred when intestinal continuity was restored. A more recent study used Ussing chambers to measure intestinal permeability in a single patient before and after fecal diversion.^33^ Diversion of the fecal stream decreased inflammation of the mucosa and normalized epithelial degeneration and mucosal permeability. The permeability changes recurred when the diversion was removed. These studies suggested a luminal factor for the onset of microscopic colitis. The gut microbiota is a logical target of investigation given the important roles of bacteria in the intact gut and the profound change when the fecal stream is diverted.

Previous studies have reported alterations in the gut microbiome associated with microscopic colitis using fecal samples and that alpha diversity of the gut microbiome was often decreased in microscopic colitis patients. Rindom Krogsgaard et al. reported stool findings from 10 patients with lymphocytic colitis, 10 with collagenous colitis and 10 healthy controls^17^ and reported that the bacterial composition of the cases differed from the controls at baseline but not after treatment with budesonide. There was decreased bacterial diversity during active disease that increased after treatment. Similarly, in a study by Hertz et al. ^18^, the stool microbiota in 15 patients with microscopic colitis was less diverse than in 21 healthy controls. With 29 cases with collagenous colitis, 29 healthy controls, 32 patients with Crohn’s disease and 32 with ulcerative colitis, Carstens et al. found loss of taxa in collagenous colitis patients with active disease and on steroid treatment that resembled the findings in the IBD patients.^14^ Morgan et al. compared 20 microscopic colitis patients to 20 healthy controls and 20 patients with functional diarrhea ^34^ and found a significant difference in alpha diversity between patients with active disease and remission. Alpha diversity was lower in the active cases compared to healthy controls and diarrhea controls, but the results were not significant.

While changes in alpha diversity were broadly consistent between studies, the microorganisms associated with microscopic colitis reported in previous studies have not been always consistent, potentially because of small sample sizes and different ways that different studies define control groups. Helal et al. cultured biopsies from 20 patients and 10 normal controls ^35^ and reported an association between *E. coli* and lymphocytic colitis. In another study, Fischer et al. examined fecal samples in 10 female patients with microscopic colitis and compared them to 7 healthy controls and observed that the patients with microscopic colitis had lower amounts of *Akkermansia*.^36^ Millien et al studied 20 cases of microscopic colitis and 20 healthy controls^19^ and reported that the cases had an increase in proinflammatory sulfur reducing bacteria with a significant decrease in the Coriobacteriaceae family that was abundant in the healthy gut. Morgan et al. found that the relative abundance of *Haemophilus parainfluenzae, Veillonella parvula*, and *Veillonella* unclassified was higher in microscopic colitis cases, and the abundance of *Alistipes putredinis* was higher in healthy controls. ^34^

In this study, we collected ascending and descending colon biopsy samples of study participants and compared the microscopic colitis cases and controls with a diarrhea history. Our approach minimizes the potential influence of diarrhea on microbial composition of fecal samples. With our relatively large sample size of 52 cases and 153 controls, we observed a decreased Shannon diversity in the descending colon microbiome associated with microscopic colitis, which verified some previous findings ^13, 14, 15^. In this study, we also reported some differences in individual taxa that are consistent with previous studies such as enrichment of Gammaproteobacteria with microscopic colitis ^37^. This consistent signal associated with Gammaproteobacteria is especially interesting because Gammaproteobacteria are known to be inflammation related ^38-40^. Also consistent with previous studies ^19, 41^, the relative abundance of family Coriobacteriaceae and genus *Collinsella* was higher in controls compared to microscopic colitis cases. Our study also had some differences from previous studies. For example, we did not observe a significant difference in mucin degrading Akkermansia that has been previously reported ^11^, which might be explained by our unique approach to defining controls.

A number of factors distinguish our study from prior studies. Our study is one of the largest studies analyzing the microbiomes of microscopic colitis patients. Our choice of controls with diarrhea is an important strength. Diarrhea can alter the gut microbiota ^20^, and thus comparing microscopic colitis cases with diarrhea to healthy controls, as done in many prior studies, risks not considering the influence of diarrhea on microbial composition. In our study, we examined adherent bacteria by obtaining biopsies from two locations in the colon rather than stool specimens as has been done in most previous studies. Adherent bacteria could interact more directly with host tissues and thus may be more relevant to the etiology of microscopic colitis than luminal bacteria obtained from feces. Our biopsies were flash frozen, compared to some other studies using formalin-fixed and paraffin-embedded tissues, which while easily obtained, can result in altered microbial composition.^19^ In our study, we also adjusted for a number of covariates including age and BMI that potentially obscure the signal of microbial associations with microscopic colitis. Our study reports some important results that are consistent with previous studies, such as the increase in Proteobacteria and decrease in alpha diversity as well as some important differences. Finally, our study finds a consistent signal in the ascending and descending colon, although the signal in the descending colon is much stronger.

There are a few limitations to our study. The ages and BMI of patients are different for cases and controls. While we controlled for age and BMI in the linear models, this may have decreased the statistical power for detecting taxa associated with cases. We used 16 rRNA sequencing techniques because low biomass makes generation of shotgun metagenome sequencing data very challenging. We therefore cannot provide information on functional genes that are related to the roles of the microbiome in microscopic colitis. Although our study is one of the largest so far, it may still lack power to detect certain differential bacteria especially in the ascending colon where the signal associated with disease does not appear to be as strong as the descending colon. Another potential limitation of our study was that we combined lymphocytic and collagenous colitis. Although these subtypes are often considered separately in the literature, similarities in risk factors, histology and response to treatment would suggest that they are subtypes of the same entity. ^42^ Our study is cross-sectional, and we therefore do not know whether the observed differences in microbial communities are a cause or a consequence of microscopic colitis. Future studies with a larger sample size and using shotgun metagenome sequencing will likely provide further insights on the potential roles of the microbial community in the development of microscopic colitis

## CONCLUSION

We analyzed the microbiome of the ascending and descending colon biopsies in 52 microscopic colitis cases and 153 controls with both cases and controls having a diarrhea history. We revealed the change of microbial communities associated with microscopic colitis that are not explained by diarrhea. The descending colon microbial alpha-diversity was significantly lower in microscopic colitis cases. Increase of inflammation related taxa and decrease of *Collinsella* was associated with microscopic colitis in the descending colon microbiome. The changes of microbial features were consistent between ascending and descending colon microbiomes but not significant in the ascending colon microbiome. The altered microbial communities in patients with microscopic colitis suggest that the patients may benefit from prevention or treatment with live biotherapeutic products.

## Abbreviations

BMI: body mass index
ASC: ascending
DES: descending
SD: standard deviation

## Acknowledgments

We thank Ms. Amber Nicole McCoy and Ms. Alondra Rodriguez for technical assistance with DNA extractions, and library preparation.

## Author Contributions

RSS, AAF, TOK, JTW and AFP contributed to the conception, design, data acquisition, analysis, and supervision of the work. SS and ICB contributed to the analysis and interpretation of the data. SS, RSS and AAF drafted the manuscript, ICB, TOK, JTW and AFP contributed to revision of the manuscript. All authors have read and approved the final manuscript.

## Data availability statement

The sequencing data analyzed in this study are available at NCBI as BioProject PRJNA768799. Analysis scripts and metadata were available at https://github.com/FodorLab/MicroscopicColitisMicrobiome.

### Patient consent for publication

Not required.

### Ethics approval

The study met the standards for the ethical treatment of participants and was approved by the Institutional Review Board of the University of North Carolina at Chapel Hill.

